# Host defense responses of CO441 and CL30 maize lines to *Fusarium graminearum* analyzed by comparative label-free quantitative proteomics

**DOI:** 10.1101/700542

**Authors:** Lana M. Reid, Illimar Altosaar

**Author notes:** Corresponding author: Illimar Altosaar, Department of Biochemistry, Microbiology and Immunology, University of Ottawa, Ottawa, Ontario, K1H 8M5, Canada.

## Abstract

Gibberella ear rot is a disease of maize associated with low yields and the production of harmful mycotoxins therein. The disease is caused by the infection of host *Zea mays* with fungal pathogen *Fusarium graminearum*. Resistant (CO441) and susceptible (CL30) inbred maize line kernels were inoculated with conidial suspensions of *F. graminearum* or water (controls). Ears of maize (cobs) from each line were harvested upon maturation and proteins were extracted from the embryo tissue of the kernels to study tissue-specific response of the host. Embryo proteins from both CO441 and CL30 lines were sequenced using mass spectrometry (LC-MS/MS) and quantified using Label Free Quantification (LFQ). Following filtering, 509 proteins were identified. These proteins were grouped into nine functional categories: *Fusarium*-derived, late embryogenesis abundant, oil-body, metabolism, stress, cellular, protein storage, metabolism, and defense. Defense proteins were up-regulated in response to infection in both CO441 and CL30 lines. Furthermore, *F. graminearum* derived proteins were only found in CL30 infected kernels suggesting that resistance may be attributed in part to the inability of *Fusarium* to establish itself in the embryo. To our knowledge this is the first successful application of LFQ mass spectrometry to the study of host-pathogen response to *F. graminearum*.

**Biological significance:** Fungal pathogen *Fusarium graminearum* is responsible for billion dollar losses in crops and contamination of global grains with harmful mycotoxins. By studying host-pathogen interactions of *Fusarium* and maize on a proteomic level with resistant and susceptible genotypes, the biological interactions occurring during infection of the maturing seed were characterized. Mature kernels of the *F. graminearum* susceptible maize line CL30 and resistant CO441 line were dissected to permit a proteomic survey of the new sporophytic generation, the embryo. Detailed knowledge of this Host-pathogen interactome will assist development of new cereal lines resistant to the rot diseases caused by *Fusarium graminearum*.

**Highlights:** Susceptible (CL30) and Resistant (CO441) lines were injected with water mock or F. graminearum LC-MS/MS of maize embryo protein extracts followed by Label Free Quantification (LFQ) permitted identification, quantification and comparison of proteomes between maize genotypes and treatments Fusarium-derived proteins were abundant only in the susceptible infected embryo Defense proteomes were up-regulated in both lines following infection nsLTP and Protease Inhibitor were significantly over-expressed in the Susceptible line after infection; chitinase and WIP1 were significantly over-expressed in the Resistant line after infection

## 1. Introduction

*Fusarium graminearum* (also referred to as *Gibberella zeae* in its sexual form) is an ascomycetic fungus and the etiological agent of crop diseases including head blight or scab disease on wheat and ear or stalk rot of maize [1]. Estimates of damage caused by *F. graminearum* in wheat alone exceeded $2.7 billion between the years 1998 and 2000, making characterization of these plant-pathogen interactions a pressing concern [2]. Furthermore, chronic exposure to *F. graminearum* in humans has been linked to neurological and immunosuppressive disorders such as toxic aleukia and Akakabi toxicosis [2,3].

The genome of the PH-1 strain of *F. graminearum* was sequenced in 2007, enabling the identification of other putative virulence determinants [4]. Among the determinants expressed by *Fusarium* and other filamentous fungi are degradative enzymes such as pectinases, amylases, and proteases, which are important for functions including nutrient acquisition and establishment in the host [5-7].

In the early stages of infection, *Fusarium* appears to colonize plant cells primarily by secreting cell-wall degrading enzymes [8]. *Fusarium* exhibits hemi-biotrophic behavior. Only after replication and intracellular growth of *F. graminearum*, cell and tissue necrosis are observed. Necrosis is thought to correlate with the release of the *Fusarium*, permitting additional infectious cycles to take place on remaining living tissue [9].

Once in the host, *Fusarium* causes biochemical changes. A major orchestrator of these biochemical changes is the trichothecene deoxynivalenol (DON). The *Tri5* gene cluster in the pathogen encodes for trichothecene synthase and disruption of this gene cluster has been linked with lower disease severity [10]. Amine inducers, low pH, reactive oxygen species (ROS) and certain phenolic acids have been recognized as inducers of DON biosynthesis [11-14]. Studies have implicated DON as a major virulence factor, however, the mechanism by which it confers virulence is unknown [15].

Infection of maize can induce various host defense mechanisms including the production of nitric oxide, ROS, phytoalexins and pathogenesis-related (PR) proteins [16,17]. Additionally, phenolic acid abundance in the cell has been linked with increased resistance to fungal disease [14]. Phenolics confer an antifungal effect and are involved in the fortification of the plant cell walls through esterification [18]. In maize and wheat, the cell walls contain an abundance of phenolic acids such as ferulic and *p*-coumaric acids. Following infection, cell wall-bound phenolic compounds increase. These hydroxycinnamic acids in particular have been linked with resistance to *Fusarium* in maize kernels [19]. The molecule *p*-hydroxybenzoic acid has been found to reduce *TRI5* expression and DON biosynthesis whereas the production of ferulic acid has been linked with activation of DON biosynthesis [14]. Therefore, resistance may stem from differences in phenolic acid production among plants.

PR proteins are defensive proteins expressed by the plant upon challenge by a pathogen and are classified into 17 families based on their role and molecular characteristics [20]. A recent study shows that a large majority of up-regulated proteins in maize following *F. graminearum* infection are PR proteins such as chitinase and thaumatin-like protein [1]. In wheat, expression of β-1, 3-glucanases (PR-2) and chitinases (PR-3) was found in greater abundance in the resistant line Arina when compared to the susceptible line, Agent [21]. These proteins confer protection by catalyzing the hydrolysis of fungal cell wall constituents, chitin and β-1, 3-glucan [22]. Further studies on the PR protein response may help identify other key defenders in the plant-*Fusarium* interactome.

The objective of the present study was to determine differences in defense protein expression between maize genotypes and in their proteomes following *F. graminearum* inoculation. It was hypothesized that defense proteins of maize embryos infected with *F. graminearum* are differentially expressed depending on their genotypes. To test this hypothesis, analysis of the maize embryo under the different experimental treatments was assessed using an aqueous protein extraction method and gel free LC-MS/MS followed by label free quantification (LFQ) proteomics.

## 2. Materials and Methods

### 2.1. Maize harvesting and kernel sampling

Experimental procedures were performed as described in [23]. *F. graminearum* resistant maize inbred line (CO441) and susceptible line (CL30) were planted on the Central Experimental Farm in Ottawa, ON, Canada. Maize kernels were inoculated with either water (control) or with *F. graminearum* conidial suspensions. Kernels were injected using a four-pin automatic kernel inoculator, 10-15 days post-silk emergence [24]. *F. graminearum* strain DAOM180378 was used and conidial suspensions were prepared at 5 × 10^5^ conidia/mL. Ears of both fungal and water treated maize were harvested at maturity and stored in brown paper seed bags at -20 °C until further analysis.

### 2.2. Protein extraction from embryo

Embryo protein extracts were obtained from each treatment (1 sample = 16 kernels). Kernels were washed with a 10% sodium hypochlorite solution for 5 min followed by five washes in ddH_2_O. Washed kernels were placed on sterile Whatman paper and left to dry overnight in the laminar flow hood. Dried kernels were incubated 15 h in 0.3% (w/v) sodium metabisulphite and 85% (v/v) lactic acid at 37 °C. Embryo tissues were dissected using a surgical scalpel. Tissue was homogenized with 20 mL of aqueous protein extraction buffer (10mM Tris-HCl pH 7.8, 0.5 mM EDTA, 10 mM KCl, 1mM PMSF (protease inhibitor) adjusted to pH 7.8) at 4 °C. Homogenates were centrifuged at 14,000 *g* for 10 min at 4 °C. 70% of the supernatant was removed and placed into fresh tubes. Pellets were re-suspended in the remaining supernatant and centrifuged a second time. Supernatants from each sample were pooled together and protein was precipitated with pre-chilled acetone (4 volumes) at -20 °C for 1 h and centrifuged at 14,000 *g* for 20 min at 4 °C. The subsequent protein pellets were washed twice with acetone and left to air dry in the laminar flow hood. Protein extracts were stored at -80 °C.

### 2.3. Quantification of protein extracts

A Bradford assay was used to determine the concentration of protein in each sample [25]. SDS-PAGE using a 15% acrylamide separating gel using Tris-glycine buffer was performed to confirm the presence of protein in each sample [26]. Ten µg of each sample was used for mass spectrometric protein sequencing and quantification by the Ottawa Institute of Systems Biology (OISB).

### 2.4. Mass Spectrometry (LC-MS/MS)

A detailed description of methods for LC-MS/MS is provided in the supplemental files (Supplemental 2). Protein extracts were dissolved in 8 M urea, reduced and alkylated. Samples were digested with trypsin and the resulting peptides were analyzed by high-performance liquid chromatography/electrospray ionization tandem mass spectrometry (HPLC-ESI-MS/MS) [27]. The acquired MS/MS spectra were searched against *Zea mays* (Uniprot taxon: 4577 and 76912) and *Gibberella zeae* PH-1 (Uniprot taxon: 229533) Uniprot protein sequence databases (version 3.85) using MaxQuant (Martinsried, Germany) with the label free quantification (LFQ) option. The false discovery rates (FDR) for both the protein and peptide levels were set to ≤1%. Quantification was performed using normalized LFQ intensity [28].

### 2.5. Data Analysis

Data from each set of biological duplicates were combined and an average of their LFQ intensities was calculated. Maize protein hits from each sample (4 samples, each with n=2) were manually assigned to a functional group based on their role in the cell, and groups were as follows: late embryogenesis abundant (LEA), protein storage, cellular, metabolism, defense, stress response, oil-body, and uncharacterized. Statistically significant changes in defense protein expression were reported when a given protein compared between two samples passed a Significance B test (at *P* < 0.05) with the MaxQuant/Perseus software (version 1.3.0.4). A full description of Significance B testing is given by Cox and Mann [28]. To better describe uncharacterized proteins, their homologs were matched using a basic local alignment search tool (BLAST) search and the NCBI protein database [29].

## 3. Results

The experimental model involved injection of maize kernels with *Fusarium* or with water (control). Visual observation of the susceptible line (CL30) indicated aggressive *Fusarium* infestation and rot (Figure 1). Neither the resistant infected and resistant water treated maize displayed any visible signs of disease.

**Figure 1:**
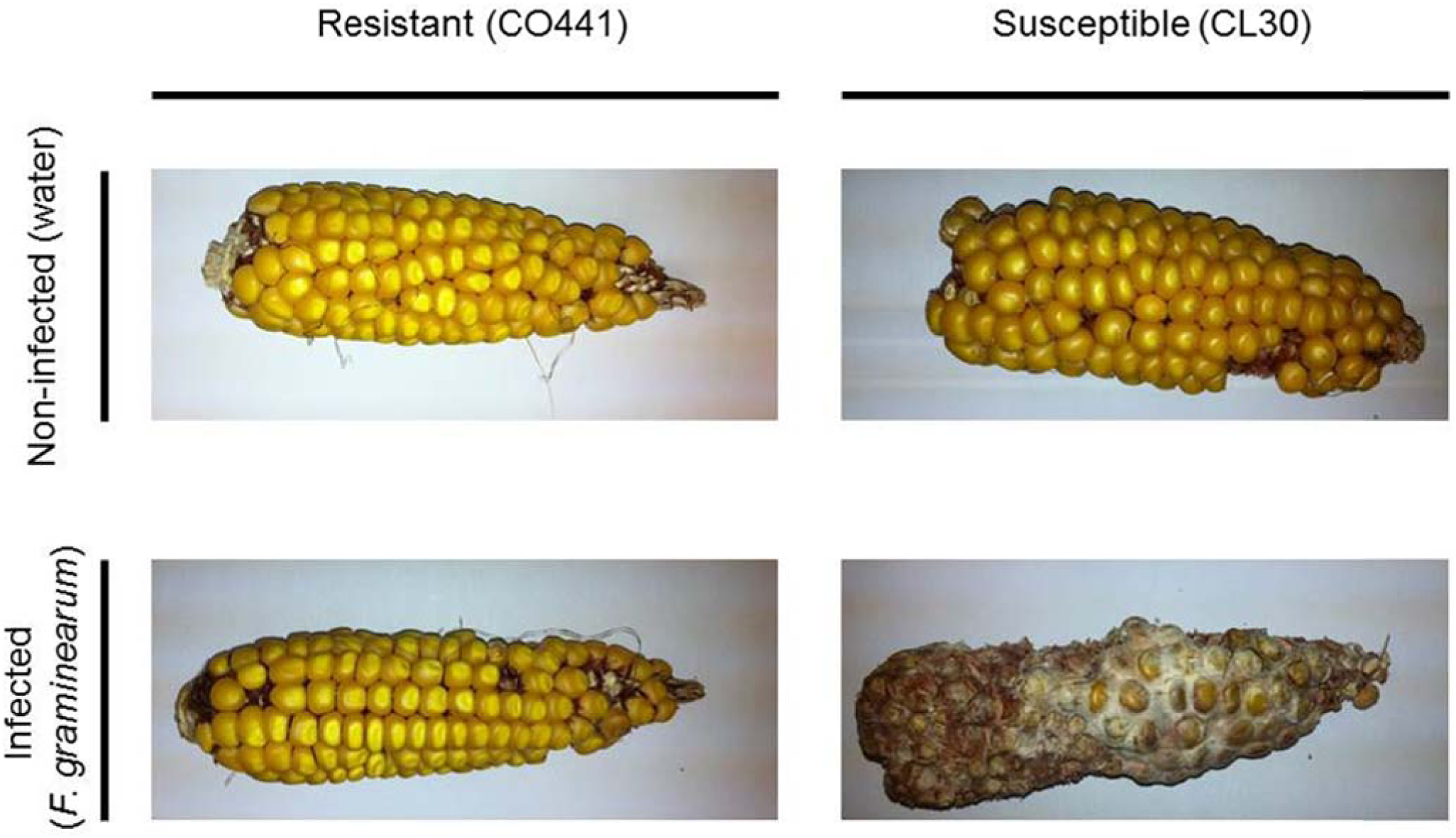
Resistant and susceptible maize injected with water or *F. graminearum*. Resistant (CO441) and susceptible (CL30) maize kernels were inoculated 10-15 days after flowering and ears were harvested at maturity at the Central Experimental Farm, Ottawa.

### 3.1. Proteins identified

Cumulatively, 951 proteins were identified following mass spectrometry. This list of protein hits was filtered for reverse hits, contaminants, posterior error probability (*PEP*) <0.05 and proteins with fewer than two peptide counts. Following filtering, the list consisted of 509 proteins in total (Supplemental 1).

### 3.2. Organismal and functional grouping of proteins

Maize protein hits were manually classified into eight groups based on their biological function in the cell: late embryogenesis abundant (LEA), oil-body, metabolism, stress, cellular, protein storage, metabolism, and defense (Figure 2).

**Figure 2:**
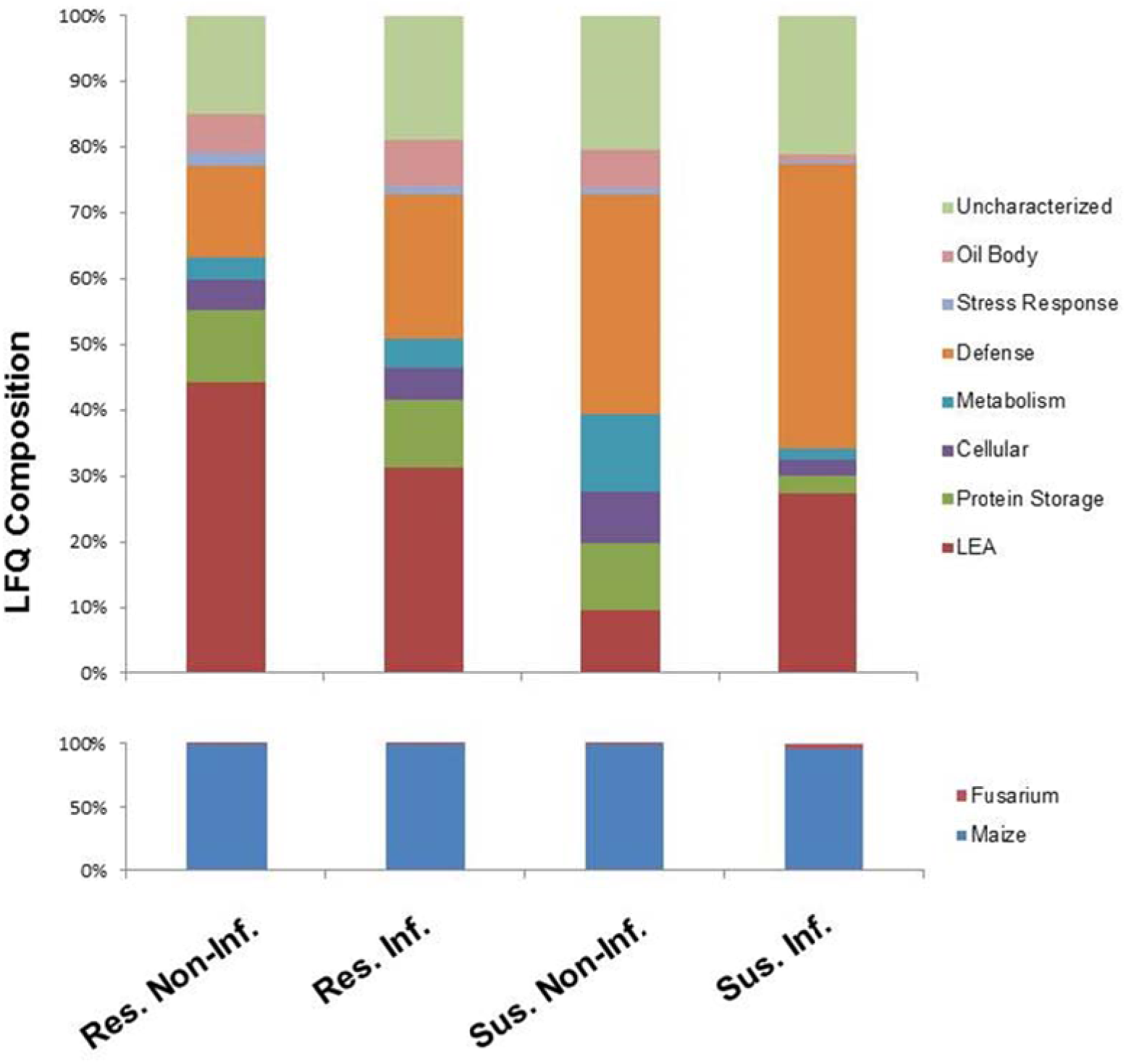
Defense protein expression responded to *F. graminearum* inoculation. **Top:** Functional grouping of maize proteins identified in the embryo. Protein IDs were manually assigned to a group depending on their biological function. Groups were as follows: Uncharacterized, Oil Body, Stress Response, Defense, Metabolism, Cellular, Protein Storage and Late Embryogenesis abundant (LEA). Label free quantification (LFQ) intensities from each functional group were summed and normalized to 100%. Samples were prepared in duplicate (n=2). **Bottom: Proportion of protein derived from maize and *F. graminearum* using LFQ intensities.** Functional grouping of proteins identified in the embryo. Protein IDs were manually assigned to a group depending on their organism of origin (*Fusarium* or maize).

LEA protein abundance was 44.2%, 31.3%, 9.64% and 27.3% of the total maize protein in resistant water-treated (R_W_), resistant *Fusarium*-treated (R_F_), susceptible water-treated (S_W_) and susceptible *Fusarium*-treated (S_F_), respectively. Following infection, the absolute value of LEA protein abundance (represented by LFQ intensity) decreased by a factor of 0.560 in the resistant line and increased by a factor of 4.06 in the susceptible line.

Storage protein abundance was 11.1%, 10.3%, 10.0% and 2.88% of the total maize protein in R_W_, R_F_, S_W_ and S_F_, respectively. Following infection, the absolute value of LEA protein abundance (represented by LFQ intensity) decreased by a factor of 0.733 in the resistant line and by a factor of 0.412 in the susceptible line.

Cellular protein abundance was 4.71%, 4.96%, 7.99% and 2.30% of the total maize protein in R_W_, R_F_, S_W_ and S_F_, respectively. Following infection, the absolute value of LEA protein abundance (represented by LFQ intensity) decreased by a factor of 0.833 in the resistant line and by a factor of 0.413 in the susceptible line.

Metabolism protein abundance was 3.21%, 4.28%, 11.7% and 1.78% of the total maize protein in R_W_, R_F_, S_W_ and S_F_, respectively. Following infection, the absolute value of LEA protein abundance (represented by LFQ intensity) increased by a factor of 1.05 in the resistant line and decreased by a factor of 0.219 in the susceptible line.

Stress response protein abundance was 1.89%, 1.27%, 1.20% and 0.513% of the total maize protein in R_W_, R_F_, S_W_ and S_F_, respectively. Following infection, the absolute value of LEA protein abundance (represented by LFQ intensity) decreased by a factor of 0.530 in the resistant line and by a factor of 0.613 in the susceptible extracts.

Oil-body protein abundance was 5.93%, 7.11%, 5.78% and 0.854% of the total maize protein in R_W_, R_F_, S_W_ and S_F_, respectively. Following infection, the absolute value of LEA protein abundance (represented by LFQ intensity) decreased by a factor of 0.947 in the resistant line and by a factor of 0.212 in the susceptible line.

Uncharacterized protein abundance was 15.0%, 18.8%, 20.3% and 21.2% of the total maize protein in R_W_, R_F_, S_W_ and S_F_, respectively. Following infection, the absolute value of LEA protein abundance (represented by LFQ intensity) decreased by a factor of 0.995 in the resistant line and increased by a factor of 1.50 in the susceptible line.

Defense protein abundance was 14.02%, 21.2%, 33.3% and 43.2% of the total maize protein in R_W_, R_F_, S_W_ and S_F_, respectively. Following infection, the absolute value of LEA protein abundance (represented by LFQ intensity) increased by a factor of 1.24 in the resistant line and by a factor of 1.86 in the susceptible extracts.

Proteins were also classified based on the organism from which they were derived (maize or *F. graminearum*) (Figure 2). Maize-derived proteins comprised the majority of proteins in the samples with *Fusarium* proteins contributing 0.0111%, 0.0124%, 0.0166% and 4.38% of the total proteins in R_W_, R_F_, S_W_ and S_F_, respectively. *Fusarium* abundance in R_F_ differed by a factor of 0.90 when compared to the non-infected control. S_F_ contained the most *Fusarium*-derived proteins, 378 times greater in comparison to the S_W_ baseline. Protein quantities used in this initial functional and organismal classification were not statistically tested for significance.

A total of 17 *Fusarium*-derived proteins were identified, all uncharacterized. A BLAST search of the amino acid sequences was used to identify proteins in other organisms that shared similar homology. The homologs with greatest similarity (sharing >50% identity) were as follows: UniProt: I1S0I4 shared 100% identity with Snodprot from *Fusarium graminearum* (Uniprot: Q5PSV7). UniProt: I1RLE0 shared 100% identity with Alkaline foam protein B found in *Fusarium culmorum* (Uniprot: A1XCJ0). UniProt: I1RQB6 shared 97.0% identity with Glyceraldehyde-3-phosphate dehydrogenase found in *Fusarium oxysporum* (Uniprot: F9FK90). UniProt: I1RL08 shared 84.0% identity with Small secreted protein from *Fusarium solani* subsp. *pisi* (Uniprot: C7Z066). UniProt: I1RC55 shared 78.0% identity with Heat shock protein 30 found in *Cylindrocarpon tonkinense* (Uniprot: C3VMP2). UniProt: I1S341 shared 78% identity with Eliciting plant response-like protein (Epl1) from *Trichoderma asperellum* (Uniprot: E3UTB4). UniProt: I1RSM6 shared 69.0% identity with Glucose repressible protein (Grg1) found in *Cordyceps militaris* strain CM01 (Uniprot: G3JKJ7) (Supplemental 1).

### 3.3. Significant changes in protein expression

Using Label Free Quantification (LFQ) intensities, a Significance B test (*P*<0.05) was performed to compare expression changes between samples with either identical genotypes (resistant/susceptible) or identical treatments (water/*Fusarium* injected). A total of 68 proteins was determined to have changed significantly in at least one two-sample comparison (Supplemental 1). Of this total, 13 defense proteins were deemed to change significantly. Defense proteins that were up-regulated following infection in the resistant line (R_F_/R_W_) were chitinase (Uniprot: B8QUX1 and Q6KBR1), basic endochitinase A (Uniprot: B6TR38), β-1, 3-glucanase (UniProt: E1AFV5), catalase (UniProt: B6UHU1) and Bowman-Birk type wound-induced proteinase inhibitor WIP1 (UniProt: B4FYK1). Proteins down-regulated between R_F_/R_W_ were non-specific lipid transfer protein (UniProt: B6T1X4, B6SP11 and B6T2T4), glutathione peroxidase (Uniprot: B6TR92) and Lipoxygenase (Uniprot: Q8W0V2) (See Table 1 for values).

In the S_F_/S_W_ and R_W_/S_W_ comparisons, glutathione peroxidase (Uniprot: B6TR92) was up-regulated. Comparing the infected resistant to the infected susceptible genotype (R_F_/S_F_), there was an up-regulation of β-1, 3-glucanase (UniProt: E1AFV5) and non-specific lipid transfer protein (Uniprot: B6SGP7). Down-regulated proteins included protease inhibitor (Uniprot: Q2XWZ0) and non-specific lipid protein (Uniprot: B6SP11). Furthermore, the protein score represented a log_2_ transformation of the posterior error probability (PEP) and was calculated by MaxQuant software. High scores indicate greater confidence of a protein being identified correctly. The highest scores were attributed to chitinases (3000 and 2860) and the lowest score was attributed to catalase (90.9) (Table 1).

## 4. Discussion

### 4.1. Susceptibility - *Fusarium* establishes itself within the host-embryo

Label Free Quantification (LFQ) permitted relative quantification of protein among all samples (Figure 2). By searching against the recently sequenced genomes of *Zea mays* and *Fusarium graminearum*, peptide hits were used to determine their corresponding protein IDs [4,30]. These Protein IDs were manually assigned to a functional group depending on their biological function in the cell. Grouping allowed for a determination of the respective effects of maize genotypes and treatment on defense protein expression. Without *F. graminearum* infection, defense protein expression changed by 0.528 fold in the resistant when compared to the susceptible proteome (R_W_/S_W_). Following infection with *F. graminearum* this value changed to 0.353 (R_F_/S_F_). Furthermore, defense protein expression increased by a factor of 1.24 in the resistant line (R_F_/R_W_) and by a factor of 1.86 in the susceptible extracts (S_F_/S_W_). This suggested that the susceptible line expressed a greater abundance of defense proteins (compared to the resistant line) regardless of treatment and that both the resistant and susceptible lines up-regulated their defense response following the fungal attack. The resistant genotype may have expressed proteins that are more effective for preventing the establishment of *F. graminearum* in the embryo. As such, more drastic changes in expression levels of the defense proteome were not required. To our knowledge, this theory has yet to be explored and consequently, further studies are needed for validation.

Proteins were also classified according to whether they were maize-derived or *Fusarium-*derived. The majority of the proteins identified (>90%) were maize-derived. A low quantity of *Fusarium* proteins were found in samples extracted from both R_W_ and S_W_. These were used as baseline values (these samples serve as negative controls). In the R_F_ protein extracts, there was very little evidence for the presence of *Fusarium* peptides remaining through the field growing season past harvest and storage time (a 0.90 fold difference in response to infection). This implies that the fungus was unable to establish itself in the embryo. In the S_F_ sample, however, there was a 378 fold increase in response to infection, implying that the fungus is able to establish itself in the embryo. From these data, we propose that resistance to *Fusarium* is due in part to the inability of the fungus to establish itself in the embryo. The expression of specific proteins may be the link between what constitutes an efficient defense response (resistant genotype) and a non-sufficient response (susceptible genotype). This finding also suggests a new research frontier, one in which proteomic sampling throughout the growing season might be able to track fungal proteins of varying persistence in the host endosperm milieu. Finally, the lower abundance of *F. graminearum* protein hits indicates that these proteins are not recalcitrant enough to remain detectable after six weeks of grain development and maturation.

### 4.2. Defense and Pathogenesis-related proteins

To achieve a more stringent picture of expression changes potentially occurring between treatments and genotypes, a Significance B test was performed. Thus, the total list of 509 proteins was reduced to 68 proteins. Using this reduced list to compare the resistant to the susceptible genotype, there was a greater number of proteins whose expression changed significantly in the resistant genotype following infection. Despite this, the overall abundance of defense proteins was greatest in the susceptible genotype following infection.

Infection of plants induces various defense mechanisms including the production of reactive oxygen species (ROS) by class III peroxidases. The generation of these ROS generates toxic environments for the pathogen [31]. Overabundances of these ROS, however, can be harmful to the host plant. To reduce this oxidative stress, glutathione peroxidases (GPxs) are expressed. GPxs prevent oxidative damage to the host membrane by lipid peroxidation repair. In transgenic tobacco, expression of *Chlamydomonas* glutathione peroxidase was found to confer greater tolerance to oxidative stress [32]. Repair from this oxidative stress is achieved through the reduction of various organic and hydro-peroxides to their respective hydroxyl compounds [32]. In the present experiment, the glutathione peroxidase level was reduced by two-fold in the resistant line following treatment with *F. graminearum* (R_F_/R_W_) (Table 1). These data suggest oxidative stress in the resistant genotype did not reach levels that would require protection of the host membrane and consequently, the expression of GPx was down-regulated.

Non-specific lipid transfer proteins (nsLTPs) are small cysteine-rich proteins involved in the shuttling of lipids and hydrophobic molecules between membranes [33]. The nsLTPs are implicated with plant defense following biotic and abiotic stresses and are able to form protective extracellular waxy and cutin polymeric layers through the deposition of hydrophobic monomers [33,34]. Furthermore, nsLTPs are able to confer antimicrobial properties *in vivo*, although this is not conserved across all nsLTPs [34]. The nsLTPs from maize leaf protein extracts were found to have an inhibitory effect on *Fusarium solani* growth [35]. In the current study, the nsLTPs increased most following infection of the susceptible genotype (S_F_/S_W_). In contrast, the resistant genotype was associated with a decrease in nsLTP levels following infection (R_F_/R_W_). This suggests that in this study, nsLTPs have a greater role in pathogen defense response in the susceptible genotype compared to the resistant. The effectiveness of nsLTPs depends both on the pathogen and the specific nsLTP expressed (there are many with different function and activity). In a previous study, *F. graminearum* was among the fungi least inhibited by wheat nsLTP [33]. There, only three out of eight nsLTPs were shown to provide at least a moderate level of antifungal activity. In the context of the current report, observations from the Sun study suggest that susceptibility is not a result of an absent defense response, but rather the result of an ineffective one. Furthermore it suggests that the resistant genotype is not as dependent on these proteins as a means of defense, relying instead on other more effective PR-proteins.

Another defense-related protein that we observed to change significantly was β-1, 3-glucanase. The enzyme β-1, 3-glucanase catalyzes the hydrolysis of fungal cell walls by cleaving β-1, 3-glucan [36]. Transgenic wheat lines expressing β-1, 3-glucanase have been linked with increased resistance to *Fusarium* infection [37]. In our maize embryos, there was no significant change in abundance of β-1, 3-glucanase between S_F_/S_W_, however, there was a 12.4 fold increase between R_F_/R_W_. These data suggest that expression of β-1, 3-glucanase is induced by *Fusarium* infection in the resistant genotype but is unaffected in the susceptible genotype.

Chitinases are major PR proteins involved in the exo or endo β-1, 4-glycosidic cleavage of N-acetylglucosamine (chitin). Fungal cell walls are comprised of chitin, therefore chitinases confer antifungal properties by disassembling the structural network of these cell walls [38,39]. When maize was challenged with *F. graminearum* we found that chitinase expression was up-regulated in the infected resistant line compared to the water treated resistant line by 8.11 fold (R_F_/R_W_). Comparing the two genotypes, chitinase (UniProt Q6JBR1) was expressed 30.68 fold greater between R_F_/S_F_, implicating it in the resistance trait. These data are consistent with another study with CO441, but one using gel spot excision. Chitinase expression was greater in the CO441 resistant genotypes in comparison to susceptible and increased further upon infection with *Fusarium* [1].

Wound-induced protease inhibitor 1 (WIP1) increased significantly following *F. graminearum* infection in the resistant line (R_F_/R_W_= 9.32). WIP1 is environmentally regulated in plant tissues and induction of *WIP1* is confined to the site of wounding [40]. Mohammadi et al, 2011 found increased WIP1 in the resistant CO441 line (both infected and non-infected samples) compared to the susceptible B73 suggesting that resistance may be due to these wound-inducible protease inhibitors [1]. In the present study, we observed an increase in expression of WIP1 in response to *Fusarium* challenge (R_F_/R_W_), however, there were no significant changes between the susceptible duplicates as a result of infection (S_F_/S_W_), reinforcing the notion that expression of this specific protease inhibitor is governed by factors other than wounding, in this case genotypic differences. These data suggest that resistance may be due to an up-regulation of WIP1 as it is important in defense against pathogen in the early stages of colonization.

Fungi secrete a variety of proteases upon infection that facilitate colonization [7]. Host plants, including maize, are known to express protease inhibitors upon fungal challenge [40,41]. In maize, protease inhibitor abundance was found to increase in resistant (CO441) and susceptible (B73) genotypes following *F. graminearum* infection and expression was overall greater in the resistant genotype compared to the susceptible, regardless of treatment (see Table 3 in [1]). The present experiment, however, produced confounding observations. Protease inhibitors were less abundant in the resistant compared to the susceptible line following infection (R_F_/S_F_= 0.00367) (Table 1). A possible explanation for this may be that the resistant line halts the infection in the early stages and as such, no additional action by the host is required to defend against *F. graminearum*. In contrast, the susceptible plant may not be able to prevent *Fusarium* from colonizing and must therefore attempt to combat fungal protease activity in real-time during the course of the disease, resulting in the up-regulation of protease inhibitors. As WIP1 was observed to be induced only in the resistant line, the susceptible line relies instead on general protease inhibitors for defense, likely at a later time than these wound-induced protease inhibitors.

### 4.3. Model of resistance – chitinase is strategic during early stages of infection

The data suggest that chitinase plays a major role in resistance: *Fusarium* is unable to establish itself in the embryo of resistant lines as chitinase destroys the fungus in early stages of infection. The chitinase response from the susceptible host is absent or inadequate due to a possible delay in salicylic acid response. As a consequence of this delayed response, *Fusarium* may be able to overpower host-defenses and infect.

The exact role of protease activity by *F. graminearum* is complex; however, it may prove important to the severity and rate of disease progression. The susceptible host genotype appears to depend primarily on nsLTP up-regulation to defend against *F. graminearum* (Table 1); however, the proteins are not sufficient or effective in combating the fungus. Consequently, the susceptible maize embryo may attempt to counteract the degradative nature of such *Fusarium*-derived proteases by expressing protease inhibitors. According to this proposed model (Figure 3) the resistant phenotype prevents early entry of *Fusarium* through increased expression of WIP1 (Table 1) and as a result, fungal establishment in the embryo is rendered unfavorable.

**Figure 3:**
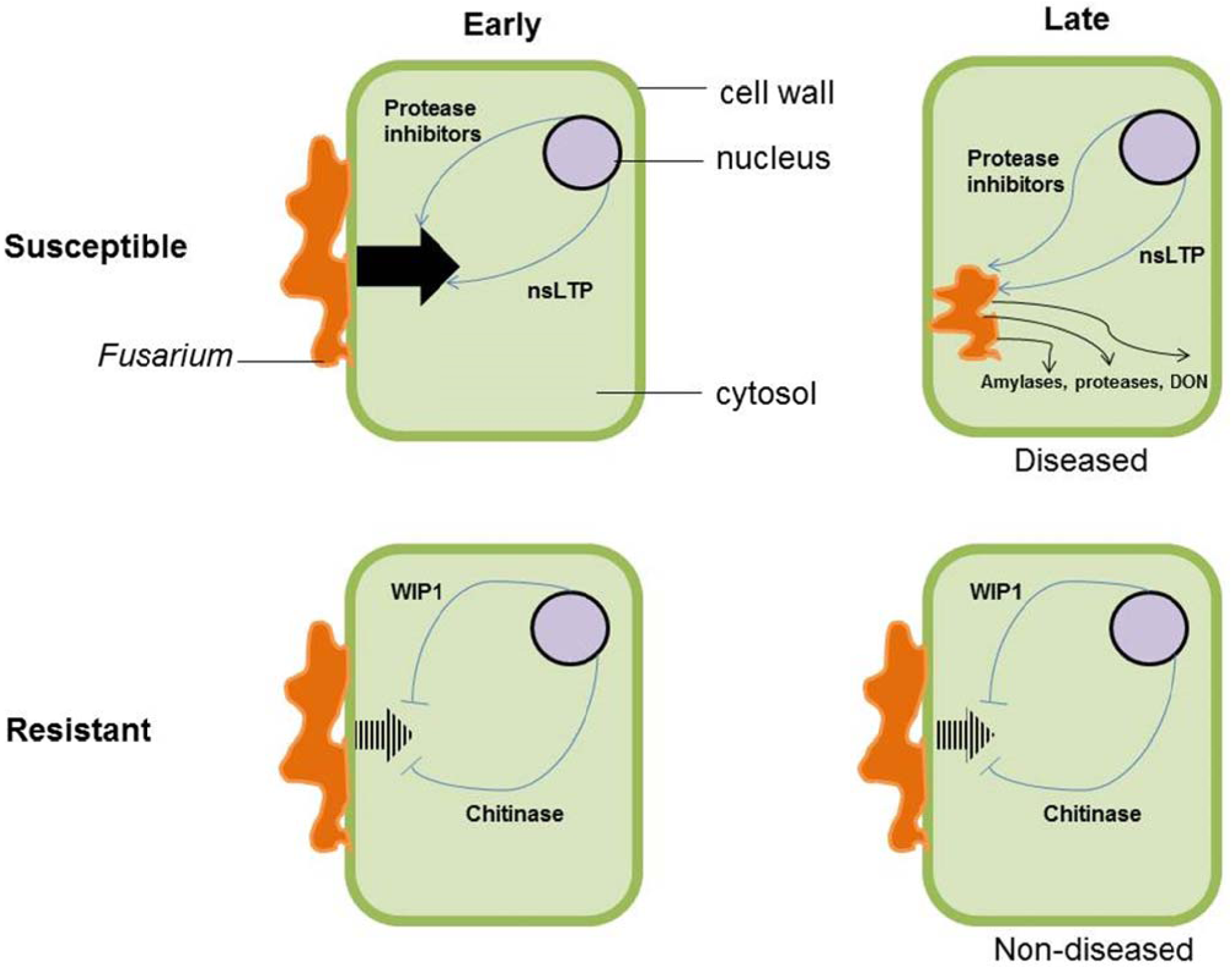
Model of *F. graminearum* pathogenesis compared between susceptible and resistant lines of maize at different stages of infection. At early stages of contact between pathogen and host, *Fusarium* (orange mycelium) is able to invade the susceptible embryo cell (big black arrow) because non-specific lipid transfer protein (nsLTP) and protease inhibitor expression is not effective (see Table 1). Inside the susceptible cell the *Fusarium* expresses compounds such as amylases, proteases and deoxynivalenol (DON) capable of leading to a diseased state. The resistant line may be able to prevent fungal entry and colonization (dashed arrow) because proteins such as chitinase and wound-inducible protease inhibitor 1 (WIP1) provide a more effective defense (Table 1).

Introduction of compounds such as salicylic acid and ethylene are known to induce increases in chitinase production and disease resistance in plants [42-44]. Salicylic acid is an essential signaling molecule for plant defense against phytopathogens and other biotic stresses [44]. Moreover, studies of *Fusarium* infection in wheat have shown that susceptibility likely arises due to the delayed activation of the salicylic defense pathway [45]. If this mechanism occurs in maize as well, it is possible that *Fusarium* colonizes the embryo of the susceptible plant due to downstream effects of this delayed salicylic acid pathway. This may result in a delayed chitinase response providing the fungus with an opportunity to colonize and penetrate more deeply into the starch rich tissue reserves of the developing seed.

In summary, given these considerations on maize response to *Fusarium* infection as measured by LFQ proteomics, LFQ has provided novel insights into the cross-talk between host and pathogen. However, a caveat to such analyses and inferences regarding plant-pathogen interactions on a proteomic level, as described in the literature to date, is the difficulty of reproducibility. Use of different inoculation methods, hosts, pathogens and protein extraction procedures not to speak of mass spectrometer differences, results in inconsistencies between experimental models and their resultant data sets. This may explain why so little overlap is found between studies, as recently underscored by Finnie et al, 2013 [46]. Using the same material and experimental techniques on the pericarp and endosperm tissue will allow for the elucidation of biochemical processes occurring from fungal attack, retaining consistency in experimental methods.

As the endosperm and pericarp are apoptotic tissues of the grain it is expected that mass spectrometric analysis will reveal relatively fewer protein hits and lower amounts of these proteins. The current results therefore predict that a greater representation of cell wall degrading enzymes and amylases will be found in the pericarp and endosperm tissues respectively, as the fungus attempts to modify its protein expression to optimize nutrient acquisition and overcome the protective outer casing of the maturing maize kernel.

## Conclusion

Cumulatively, 509 proteins were identified in the eight maize embryo samples and functionally classified in accordance to their biological function. The defense group in resistant and susceptible maize increased following infection by *F. graminearum*. Interestingly, the stress response and uncharacterized groups showed least change following fungal challenge in the embryo. Furthermore, these data indicated *F. graminearum* is able to establish itself in the susceptible embryo but not in the resistant. When testing for significantly expressed proteins between samples, up-regulation of chitinases and non-specific lipid transfer proteins was observed following infection in the resistant and susceptible genotypes, respectively. Therefore, it was proposed that the diseased phenotype is directly linked to an inadequate defense response. A successful defense response is dependent on the ability of disarming *F. graminearum* before the fungus establishes itself in the host.

## Acknowledgements

Project was funded by Natural Sciences and Engineering Research Council of Canada (NSERC). We would also like to thank Dr. Zhibin Ning (OISB), Dr. Adam Koziol (CFIA), and Jason El Bilali for their technical assistance in this project; and we are very grateful to Dr. Kin Chan for stimulating guidance and helpful discussions.

