## Supplemental 1 for "Host defense responses of CO441 and CL30 maize lines to *Fusarium graminearum* analyzed by comparative label-free quantitative proteomics"

| Uniprot ID | Fasta headers | Proteins | Peptides | Unique peptides | Sequence coverage [%] | PEP | LFQ Resistant (CO441) Infected | LFQ Resistant (CO441) Infected- 2 | LFQ Susceptible (CL30) Infected | LFQ Susceptible (CL30) Infected- 2 | LFQ Resistant (CO441) Non-Infected | LFQ Resistant (CO441) Non-Infected- 2 | LFQ Susceptible (CL30) Non-Infected | LFQ Susceptible (CL30) Non-Infected- 2 | P-value ($R_F/R_W$) | P-value ($R_F/S_F$) | P-value ($S_F/S_W$) | P-value ($R_W/S_W$) | $R_F/R_W$ | $R_F/S_F$ | $S_F/S_W$ | $R_W/S_W$ | $R_F/R_W$ | $S_F/S_W$ | $R_W/S_W$ | $R_F/S_F$ |
|---|---|---|---|---|---|---|---|---|---|---|---|---|---|---|---|---|---|---|---|---|---|---|---|---|---|---|
| B8QUX1 | ZEAMP Chitinase | 1 | 25 | 1 | 77.6 | 1.83E-300 | 0 | 1.94E+07 | 0 | 0 | 0 | 3363700 | 0 | 0 | 0.038665575 | 1 | 1 | 1 | + | | | | 5.75408 | | | |
| Q6JBR1 | ZEAMP Chitinase | 1 | 20 | 3 | 68.3 | 1.37E-286 | 0 | 6.44E+07 | 2099800 | 0 | 2817600 | 4969900 | 0 | 0 | 0.013085794 | 0.0125053 | 1 | 1 | + | + | | | 8.271717 | | | 30.67721 |
| E1AFV5 | MAIZE Beta-1,3-glucanase | 3 | 20 | 20 | 69.3 | 1.45E-237 | 4.77E+08 | 9.36E+08 | 6.80E+07 | 7.68E+07 | 1.69E+07 | 9.71E+07 | 2.27E+07 | 1.54E+07 | 0.003249883 | 0.12232763 | 0.509062774 | 0.627158356 | + | | | | 12.40162 | | | |
| B6SGP7 | MAIZE Non-specific lipid-transfer proteir | 3 | 8 | 1 | 56.2 | 2.15E-203 | 0 | 2.55E+08 | 0 | 4043300 | 0 | 2.24E+08 | 0 | 3.05E+07 | 0.823149666 | 0.00195505 | 0.340465096 | 0.360151857 | | + | | | | | | 63.08956 |
| B4FYK1 | MAIZE Bowman-Birk type wound-induced proteinase inhibitor WIP1 | 1 | 11 | 3 | 62.9 | 2.57E-100 | 8.68E+07 | 2.14E+08 | 2.66E+08 | 7.27E+07 | 0 | 3.22E+07 | 2394200 | 6488400 | 0.008857125 | 0.7721189 | 0.095296745 | 0.563167927 | + | | | | 9.320687 | | | |
| Q2XWZ0 | ZEAMP Protease inhibitoı | 4 | 6 | 1 | 90.4 | 1.15E-58 | 0 | 1146600 | 1.66E+08 | 1.46E+08 | 567800 | 564140 | 5.82E+07 | 3.99E+07 | 0.930097814 | 0.00120242 | 0.559914052 | 0.05990023 | | + | | | | | | 0.003666 |
| Q8W0V2 | MAIZE Lipoxygenase | 4 | 7 | 7 | 12.5 | 2.80E-34 | 0 | 463030 | 0 | 0 | 1.13E+07 | 0 | 0 | 0 | 0.002483934 | 1 | 1 | 1 | + | | | | 0.040889 | | | |
| B6T1X4 | MAIZE Non-specific lipid-transfer proteir | 6 | 4 | 2 | 27.1 | 8.12E-30 | 6498500 | 9990700 | 5.83E+07 | 6.79E+07 | 1.90E+08 | 2.29E+07 | 2819400 | 5311500 | 0.015943989 | 0.18732283 | 0.202304384 | 0.126906526 | + | | | | 0.07731 | | | |
| B6SP11 | MAIZE Nonspecific lipid-transfer proteir | 1 | 7 | 7 | 38.8 | 2.58E-23 | 0 | 2.79E+07 | 3.38E+08 | 1.50E+09 | 2.15E+08 | 5.44E+07 | 8459400 | 3.34E+07 | 0.033260092 | 0.01331691 | 0.083609537 | 0.393339657 | + | + | | | 0.103449 | | | 0.015176 |
| B4FB54 | MAIZE Non-specific lipid-transfer proteir | 2 | 5 | 5 | 50.4 | 1.42E-17 | 1.65E+07 | 1.53E+07 | 4.88E+08 | 3.77E+07 | 1.37E+08 | 5106600 | 0 | 2184100 | 0.165272408 | 0.08311596 | 0.013371327 | 0.049618306 | | | + | + | | 240.6895 | 65.24271 | |
| B6TR92 | MAIZE Glutathione peroxidase | 4 | 5 | 2 | 17.9 | 4.20E-17 | 3340400 | 2892000 | 0 | 2340300 | 1.12E+08 | 1.12E+07 | 1.37E+07 | 0 | 0.004788108 | 0.64140134 | 0.41127128 | 0.310395501 | + | | | | 0.050525 | | | |
| B8QWQ8 | ZEAMP Pathogenesis-related maize seed protei | 18 | 5 | 5 | 33.5 | 5.26E-13 | 1.74E+07 | 3.85E+07 | 3.53E+07 | 6757100 | 2560500 | 0 | 1.48E+07 | 0 | 0.000334005 | 0.94226184 | 0.59431597 | 0.446845334 | + | | | | 21.82582 | | | |
| B6T2T4 | MAIZE Nonspecific lipid-transfer proteir | 1 | 4 | 1 | 42.7 | 1.13E-12 | 1.08E+07 | 3892900 | 4.93E+07 | 9.62E+07 | 1.63E+08 | 5.29E+07 | 5750400 | 4166900 | 0.011209444 | 0.1445923 | 0.211121647 | 0.150782494 | + | | | | 0.06786 | | | |
| B6TR38 | MAIZE Basic endochitinase A | 1 | 3 | 3 | 18.9 | 2.34E-12 | 6.63E+07 | 3.36E+07 | 0 | 0 | 3464700 | 6222300 | 700090 | 0 | 0.006268228 | 1 | 1 | 0.222556216 | + | | | | 10.31816 | | | |
| B6UHU1 | MAIZE Catalase | 4 | 4 | 4 | 10 | 8.10E-10 | 1.80E+07 | 0 | 0 | 0 | 2840900 | 0 | 0 | 0 | 0.02933627 | 1 | 1 | 1 | + | | | | 6.343764 | | | |
