## Supplemental 2 for "Host defense responses of CO441 and CL30 maize lines to *Fusarium graminearum* analyzed by comparative label-free quantitative proteomics"

**Mass Spectrometry Protocol**

Urea (8M) in 50mM ammonium bi-carbonate (ABC) was used to reconstitute proteins. Reduction and alkylation were done by adding dithiothreitol (DTT) to a final concentration of 20 mM at 56°C for 30min followed by 40mM indole-3-acetic acid (IAA) at room temperature. The solution was then diluted 5 times by adding 50mM ABC. Trypsin was added to achieve a protein-enzyme ratio of 50:1. Digestion was performed at 37°C overnight, with continuous shaking. Digested peptides were then desalted on Sep-Pak column (Waters) and dried down in SpeedVac (ThermoFisher Scientific, San Jose, CA). Dried peptides were reconstituted in 0.5% (v/v) FA.

All MS analyses were done by HPLC-ESI-MS/MS. The system consisted of an Agilent 1100 micro-HPLC system (Agilent Technologies, Santa Clara, CA, USA) coupled with an LTQ-Orbitrap mass spectrometer (ThermoFisher Scientific, San Jose, CA) equipped with a nano-electrospray interface operated in positive ion mode. The mobile phases consisted of 0.1%(v/v) formic acid (FA) in water as buffer A and 0.1% (v/v) FA in acetonitrile as buffer B. Peptide separation was performed on a 75µm × 100 mm analytical column packed in-house with reverse phase Magic C18AQ resins (1.9µm; 120-Å pore size; Dr. Maisch GmbH, Ammerbuch, Germany). Briefly, the sample was loaded on the column using 98% buffer A at a flow rate of 1.5µL/min for 15min.  Then, a gradient from 5% to 35% buffer B (20~50% for APols depletion test) was performed in 120 min at a flow rate of ~300nL/min obtained from splitting a 20 µL/min through a restrictor. The MS method consisted of one full MS scan from 300 to 1700 m/z followed by data-dependent MS/MS scan of the five most intense ions, a dynamic exclusion repeat count of 2, and a repeat duration of 90 sec. As well, for the experiments on the Orbitrap MS the full MS was performed in the Orbitrap analyzer with R = 60,000 defined at m/z 400, while the MS/MS analyses were performed in the LTQ. To improve the mass accuracy, all the measurements in Orbitrap mass analyzer were performed with internal recalibration (“Lock Mass”). On the Orbitrap, the charge state rejection function was enabled, with single and “unassigned” charge states rejected.

The raw files generated by the LTQ-Orbitrap were processed and analyzed using MaxQuant, Version 1.2.2.5 using the Uniprot protein FASTA database (2012, July version), including commonly observed contaminants. The following parameters were used: cysteine carbamidomethylation was selected as fixed modification; methionine oxidation, protein N-terminal acetylation, and enzyme specificity was set to trypsin. Up to two missing cleavages of trypsin were allowed. Precursor ion mass tolerances were 7 ppm, and fragment ion mass tolerance was 0.8 Da for MS/MS spectra. If the identified peptide sequences from one protein were equal to or contained within another protein's peptide set, then the proteins were grouped together and reported as one protein group. The false discovery rate (FDR) for peptide and protein was set at 1% and a minimum length of six amino acids was used for peptide identification. Quantification was performed using normalized LFQ intensity (Ning, Z; OISB, personal communication).
